# Using genetic path analysis to control for pleiotropy in a Mendelian randomization study

**DOI:** 10.1101/650192

**Authors:** Frank D Mann, Andrey A Shabalin, Anna R Docherty, Robert F Krueger

## Abstract

**Background:** When a randomized experimental study is not possible, Mendelian randomization studies use genetic variants or polygenic scores as instrumental variables to control for gene-environment correlation while estimating the association between an exposure and outcome. Polygenic scores have become increasingly potent predictors of their respective phenotypes, satisfying the relevance criteria of an instrumental variable. Evidence for pleiotropy, however, casts doubt on whether the exclusion criteria of an instrumental variable is likely to hold for polygenic scores of complex phenotypes, and a number of methods have been developed to adjust for pleiotropy in Mendelian randomization studies.

**Method:** Using multiple polygenic scores and path analysis we implement an extension of genetic instrumental variable regression, genetic path analysis, and use it to test whether educational attainment is associated with two health-related outcomes in adulthood, body mass index and smoking initiation, while estimating and controlling for both gene-environment correlations and pleiotropy.

**Results:** Genetic path analysis provides compelling evidence for a complex set of gene-environment transactions that undergird the relations between educational attainment and health-related outcomes in adulthood. Importantly, results are consistent with education having a protective effect on body mass index and smoking initiation, even after controlling for gene-environment correlations and pleiotropy.

**Conclusions:** The proposed method is capable of addressing the exclusion criteria for a sound instrumental variable and, consequently, has the potential to help advance Mendelian randomization studies of complex phenotypes.

## Using genetic path analysis to control for pleiotropy in a Mendelian randomization study

Mendelian randomization refers to the random assortment of genes that are given to children by their parents at the time of conception (1). This results in distributions of genes that are independent of many factors that often confound associations documented in observational studies (2,3). Mendelian randomization studies use genetic variants or genetic propensity scores, also called polygenic risk scores, as instrumental variables to control for gene-environment correlation when testing a putatively casual relation between an exposure and outcome. The present study focuses on the use of polygenic scores to conduct Mendelian randomization studies, with emphasis placed on reviewing whether polygenic scores meet the criteria for a sound instrumental variable. We then present an extension of genetic instrumental variable regression (4), genetic path analysis, to help overcome a limitation inherent to Mendelian randomization studies of complex phenotypes, specifically the high potential for pleiotropic effects on the exposure and outcome of interest. Using genetic path analysis, we then test whether educational attainment is associated with body mass index (BMI) and smoking initiation in a large sample of adults while estimating both gene-environment correlation and pleiotropy.

Gene-environment correlation refers to the non-random assortment of individuals into environments based on their genotype and is behaviorally manifest by individuals actively shaping and responding to their environments based, at least partly, on their heritable characteristics (5,6). This process results in heritable variation in measures of the environment (7), which, in turn, are thought to further reinforce the expression of relevant phenotypes. Importantly, without accounting for heritable variation in environmental exposures, one cannot know whether an association between an exposure and outcome reflects a true causal relation or, on the other hand, a niche-picking process (8). Auspiciously, as summary data from genome-wide association studies (GWASs) becomes readily available, it has become increasingly popular to use polygenic scores as instrumental variables for inferring causation in non-experimental studies (a.k.a. Mendelian randomization studies).

A polygenic score may be defined “as a single value estimate of an individual’s propensity to a phenotype” calculated by computing the sum of risk alleles corresponding to a phenotype in each individual, weighted by their effect size estimate from the most powerful GWAS on the phenotype (9). A polygenic score is typically calculated as PGS_*k*_ = ∑_*i*_ *β*_*i*_ SNP_*ik*_, where PGS for individual *k* in the target sample is calculated by the summation of each SNP (measured for both the person *k* and passing a set association threshold in the discovery GWAS) multiplied by the effect size, *β*, of that SNP in the discovery GWAS. Thus, polygenic scores provide an index of an individual’s genetic propensity for a given phenotype, or “an individual-level genome-wide genetic proxy” (9). Although polygenic scores may be used for a variety of purposes, a lot of emphasis has been placed on using polygenic scores as instrumental variables. However, as noted and addressed by others, it is not clear that polygenic scores meet the necessary criteria for a sound instrumental variable (4,10,11).

There are three criteria for a sound instrumental variable (12). First, sometimes called the relevance criteria, the instrument must be related to the environmental exposure. Second, according to the exclusion criteria, conditional on the relation between the exposure and outcome, there is no direct relation between the instrument and the outcome. Put differently, any relation between the instrument and outcome must be fully accounted for by its relation to the exposure. Third, the instrument should not be related to any unmeasured confounders. Note, however, that this third criteria, sometimes called the independence criteria, is not unique to using polygenic scores as instrumental variables, or instrumental variable analysis more generally, as this concern applies to all non-experimental studies for which an unmeasured confounder exists.

Nevertheless, as the size of GWASs continue to grow, polygenic scores have become increasingly potent predictors of their respective phenotypes, satisfying the relevance criteria. On the other hand, genetic correlations across related and seemingly unrelated phenotypes provides evidence for pleiotropic effects. This suggests that polygenic scores likely violate the exclusion criteria, and, therefore, casts doubt on their use as instrumental variables. In response to this concern, a number of methods have been developed to help correct for the presence of pleiotropy. For example, statistical techniques have been developed that are more robust to pleiotropic effects violating the exclusion criteria, including Egger regression (10) and summary data-based multiple regression (13), as well as pleiotropy-robust Mendelian randomization (11) and genetic instrumental variable regression (4). The present study intends to contribute to this body of work by integrating two existing methods, genetic instrumental variable regression and path analysis, to estimate and control for pleiotropy in a Mendelian randomization study using multiple polygenic scores.

In a traditional Mendelian randomization study, two regressions are estimated simultaneously: the environmental exposure is regressed on the genetic instrument, and the outcome of interest is regressed on the environmental exposure. Unfortunately, due to pleiotropic effects, the association between the genetic instrument and the outcome is not fully mediated by the association between the genetic instrument and the exposure. Put differently, conditional on the association between the exposure and outcome, the genetic instrument is often predictive of both the environmental exposure *and* outcome, violating the exclusion criteria of a sound instrumental variable. However, as summary statistics from GWASs become available for a number of social, relational, and environmental exposures, in addition to outcomes of clinical and epidemiological interest, a path analysis using polygenic scores for an exposure *and* outcome can provide an estimate and control for pleiotropy when conducting a Mendelian randomization study.

An example of a path analysis using multiple polygenic scores is depicted in Figure 1. Similar to a traditional instrumental variable analysis, an environment or exposure (E) is regressed on a genetic instrument (PRS_E_), which estimates and controls for gene-environment correlation. An outcome (Y) is then regressed on the exposure (E) free of genetic confounds that result from active and evocative gene-environment correlations. To estimate and control for the potential pleiotropic effects of the genetic instrument, a second genetic instrument is introduced (PRS_Y_), which provides an index of polygenic liability for the outcome (Y). The correlation between the genetic instrument for the exposure (PRS_E_) and the genic instrument for the outcome (PRS_Y_) can be freely estimated, while simultaneously regressing the exposure (E) and outcome (Y) on the genetic instrument for the outcome (PRS_Y_). These parameters provide a test and simultaneous control for pleiotropy, while also estimating and controlling for additional gene-environment correlations that may not have been captured by the first genetic instrument. The correlation between the two genetic instruments sheds light on whether genetic liability for the exposure has pleiotropic effects on the outcome, and the regression of the outcome on its polygenic score provides a statistical control for pleiotropy. Finally, the regression of the exposure on the genetic instrument for the outcome tests for potential gene-environment correlations not fully accounted for by the genetic instrument for the exposure. Hereinafter, we provide a demonstration of this method focusing on the relationship between education and two important health-related outcomes: body mass index (BMI) and smoking initiation.

**Figure 1.**
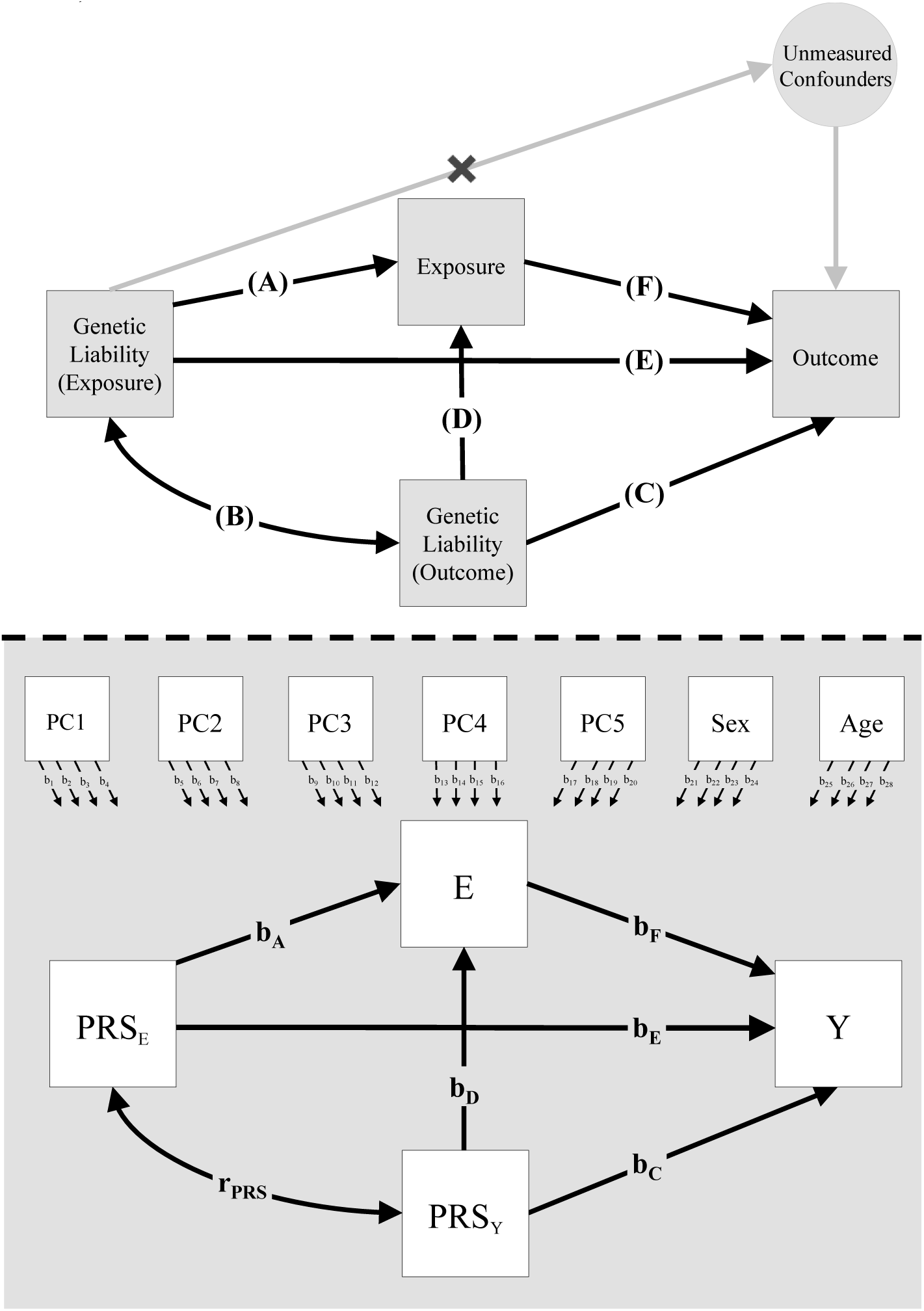
Conceptual Diagram (Top Panel) and Path Diagram (Bottom Panel) of a Genetic Path Analysis Using Multiple Polygenic Scores. **Notes.** *Top panel*: (A) test of gene-environment correlation. (B) test of pleiotropy. (C) statistical control for pleiotroy. (D) additional test for gene-environment correlation. (E) test of statistical control for pleiotropy. (F) test of quasi-causal effect of the exposure. The “X” on the pathway to unmeasured confounders reflects the independence criteria of a sound instrument. *Bottom panel*: PRS = polygenic score. E = measure of exposure. Y = measure of outcome. PC = principal component. b_1_ – b_28_ = effects of covariates on focal variables truncated to ease presentation. b_A_ = regression of exposure on polygenic risk for the exposure. r_PRS_ = correlation between polygenic risk for the exposure and polygenic risk for the outcome. b_C_ = regression of the outcome on polygenic risk for the outcome. b_D_ = regression of the exposure on polygenic risk for the outcome. b_E_ = regression of the outcome on polygenic risk for the exposure. b_F_ = regression of the outcome on the exposure.

## Method

### Sample

The present study analyses data from the Study of Midlife Development in the United States (MIDUS). (14). Data was prepared for analyses with R version 3.5.2. Data was imported into R using the ‘Hmisc’ package (15), preprocessed, and then exported from R using the ‘MplusAutomation’ package version 0.7.1 (16). Phenotype data and study materials are available on a permanent third-party archive, the 71 Inter-University Consortium for Political and Social Research (ICPSR). Additional information regarding participant recruitment, compensation, and data collection can be found elsewhere (14). Only data from participants who were genotyped and predominantly of European ancestry were included in the present study (N = 1296). The average age of participants was approximately 54 years (median = 54 years, SD = 12.46 years, min. = 25 years, max. = 84 years), and approximately 51% of the sample was female (∼ 49% male). There was considerable variation in highest level of education completed by participants (see Table 1).

**Table 1.** Highest Level of Education Completed by Participants.

|  | Order-Categorical Response |  |  |  |  |  |  |  |  |  |  |  |  |
| --- | --- | --- | --- | --- | --- | --- | --- | --- | --- | --- | --- | --- | --- |
|  | (1) | (2) | (3) | (4) | (5) | (6) | (7) | (8) | (9) | (10) | (11) | (12) | NA |
| Frequency | 1 | 6 | 26 | 10 | 199 | 195 | 59 | 110 | 325 | 54 | 232 | 70 | 6 |
| Percent | < 1% | < 1% | ~2% | < 1% | ~15% | ~15% | ~5% | ~9% | ~25% | ~4% | ~18% | ~5% | < 1% |

**Notes.** (1) = No school/some grade school (grades 1-6). (2) = Eighth grade/junior high school (grades 7-8). (3) = Some high school (grades 9-12, No Diploma or GED). (4) = GED (general education diploma). (5) = Graduated from high school. (6) = One to two years of college, no degree yet. (7) = Three or four years of college, no degree yet. (8) = Graduated from two years of college, vocational school, or obtained assoc. degree. (9) = Graduated from a four- or five-year college or obtained a bachelor's degree. (10) = Attended some graduate school, no graduate degree yet. (11) = Master's degree. (12) = PH.D., ED.D., MD, DDS, LLB, LLD, JD, etc. NA = missing values.

### Measures

The present study includes six focal constructs. Educational attainment was measured using self-reports of the highest level of education completed by participants, rated on an ordered-categorical scale. BMI was calculated based on participants height and weight (mean = 28.79, median = 27.89, SD = 6.19, min. = 17.08, max. = 77.58). There was a single outlier on BMI that was more than 5 standard deviations above than the mean; Effect sizes are similar, and results of null hypothesis significance tests remain unchanged after excluding this observation. Smoking initiation was measured by asking participants whether they were ever a smoker or currently a smoker of cigarettes (No = 59%, Yes = 41%). Polygenic scores for educational attainment, BMI, and smoking initiation were calculated using summary statistics from recent GWASs for each variable (17-19).

### Data Analytic Procedures

Path analysis was conducted in Mplus version 8.1 (20), and missing data were handled using full-information maximum likelihood (21). Because a subset of sibling-and twin-pairs are included in the current sample (N_pairs_ = 96), a family identification number was specified as a cluster variable in path models to implement a Huber-White sandwich estimator, which adjusts the standard errors of path coefficients for the non-independence of observations that results from a subset of participants being nested within the same family. Age (centered at 54 years) and biological sex (coded female = 0, male = 1) were included as exogenous covariates of all focal study variables, in addition to the first five genomic principal component scores. Thus, we report results from fully-saturated models (i.e. model degrees of freedom = 0). As the variance of certain PC scores approached zero, all PC scores were increased by a factor of 100 to avoid a singular observed covariance matrix of independent variables. BMI and smoking initiation are continuous and binary outcomes, consequently, the estimated pathways to BMI and smoking initiation can be interpreted as linear and Poisson regression coefficients, with linear coefficients standardized and Poisson coefficients exponentiated (i.e. reported as risk ratios). 99% biased-corrected bootstrapped confidence intervals are reported below their respective point estimates. Polygenic scores, self-reports of educational attainment, and BMI were standardized before fitting path models (M = 0, SD = 1).

## Results

Results for educational attainment and BMI are reported in Figure 2. Results for educational attainment and smoking initiation are reported in Figure 3. The effects of exogenous covariates are reported in Table 2. Several results are noteworthy. In both models, polygenic propensity for educational attainment was associated with educational attainment (β = .27, SE = .03, p < .001), providing evidence for gene-environment correlation. Providing evidence for pleiotropic effects, polygenic propensity for educational attainment was negatively correlated with polygenic risk for high BMI (r = -.17, SE = .03, p < .001) and polygenic risk for smoking initiation (r = -.16, SE = .03, p < .001). Providing a partial control for pleiotropic effects, polygenic risk for high BMI was associated with BMI (β = .23, SE = .03, p < .001), and polygenic risk for smoking initiation was associated with smoking initiation (RR = 1.16, SE = .04, p < .001). After accounting for these associations, the pathway from polygenic propensity for educational attainment to BMI approached zero (β = -.01, SE = .03, p = .691), as did the pathway from polygenic propensity for educational attainment to smoking initiation (RR = 0.97, SE = .04, p = .462. These estimates suggest that the regression of BMI and smoking initiation on their respective polygenic scores provided an adequate statistical control for the pleiotropic effects of polygenic risk for educational attainment.

**Table 2.** Effects of Exogenous Covariates on Focal Study Variables.

| Outcome = BMI |  |  |  |  |  |  |  |  |  |  |  |  |
| --- | --- | --- | --- | --- | --- | --- | --- | --- | --- | --- | --- | --- |
|  | PRSE |  |  | PRSY |  |  | Exposure (E) |  |  | Outcome (Y) |  |  |
|  | *b* | *SE* | *p* | *b* | *SE* | *p* | *b* | *SE* | *p* | *b* | *SE* | *p* |
| Age | .00 | .00 | .617 | -.01 | .00 | .026 | -.01 | .00 | .005 | .00 | .00 | .892 |
| Sex | .09 | .06 | .120 | -.09 | .05 | .121 | .16 | .06 | .006 | .19 | .06 | .001 |
| PC1 | -.69 | .78 | .376 | .55 | .73 | .448 | .23 | .71 | .746 | .06 | .60 | .923 |
| PC2 | -.72 | .76 | .345 | -2.44 | .69 | < .001 | .29 | .66 | .665 | .65 | .66 | .322 |
| PC3 | -.11 | .12 | .364 | .17 | .12 | .153 | .09 | .09 | .346 | .25 | .09 | .008 |
| PC4 | .00 | .15 | .467 | -.11 | .15 | .467 | -.10 | .14 | .438 | -.03 | .14 | .837 |
| PC5 | .01 | .03 | .799 | -.15 | .03 | < .001 | -.04 | .03 | .123 | .04 | .023 | .074 |

| Outcome = Smoking |  |  |  |  |  |  |  |  |  |  |  |  |
| --- | --- | --- | --- | --- | --- | --- | --- | --- | --- | --- | --- | --- |
|  | PRSE |  |  | PRSY |  |  | Exposure (E) |  |  | Outcome (Y) |  |  |
|  | *b* | *SE* | *p* | *b* | *SE* | *p* | *b* | *SE* | *p* | *b* | *SE* | *p* |
| Age | .00 | .00 | .617 | .00 | .00 | .070 | -.01 | .00 | .010 | .01 | .00 | < .001 |
| Sex | .09 | .06 | .120 | -.05 | .06 | .360 | .15 | .06 | .007 | .21 | .07 | .003 |
| PC1 | -.69 | .78 | .375 | .70 | .75 | .351 | .26 | .70 | .709 | .99 | .79 | .210 |
| PC2 | -.72 | .76 | .342 | .19 | .68 | .780 | .38 | .65 | .556 | 1.21 | .79 | .124 |
| PC3 | -.11 | .14 | .452 | .03 | .11 | .759 | .09 | .09 | .366 | .10 | .13 | .438 |
| PC4 | .00 | .15 | .999 | -.11 | .14 | .452 | -.10 | .14 | .464 | -.08 | .17 | .654 |
| PC5 | .01 | .03 | .799 | -.04 | .03 | .206 | -.04 | .03 | .137 | -.02 | .03 | .536 |

**Notes.** *b* = multiple regression coefficient. *SE* = standard error. *p* = probability of the observed data if the null hypothesis is true (i.e. *b* = 0).

**Figure 2.**
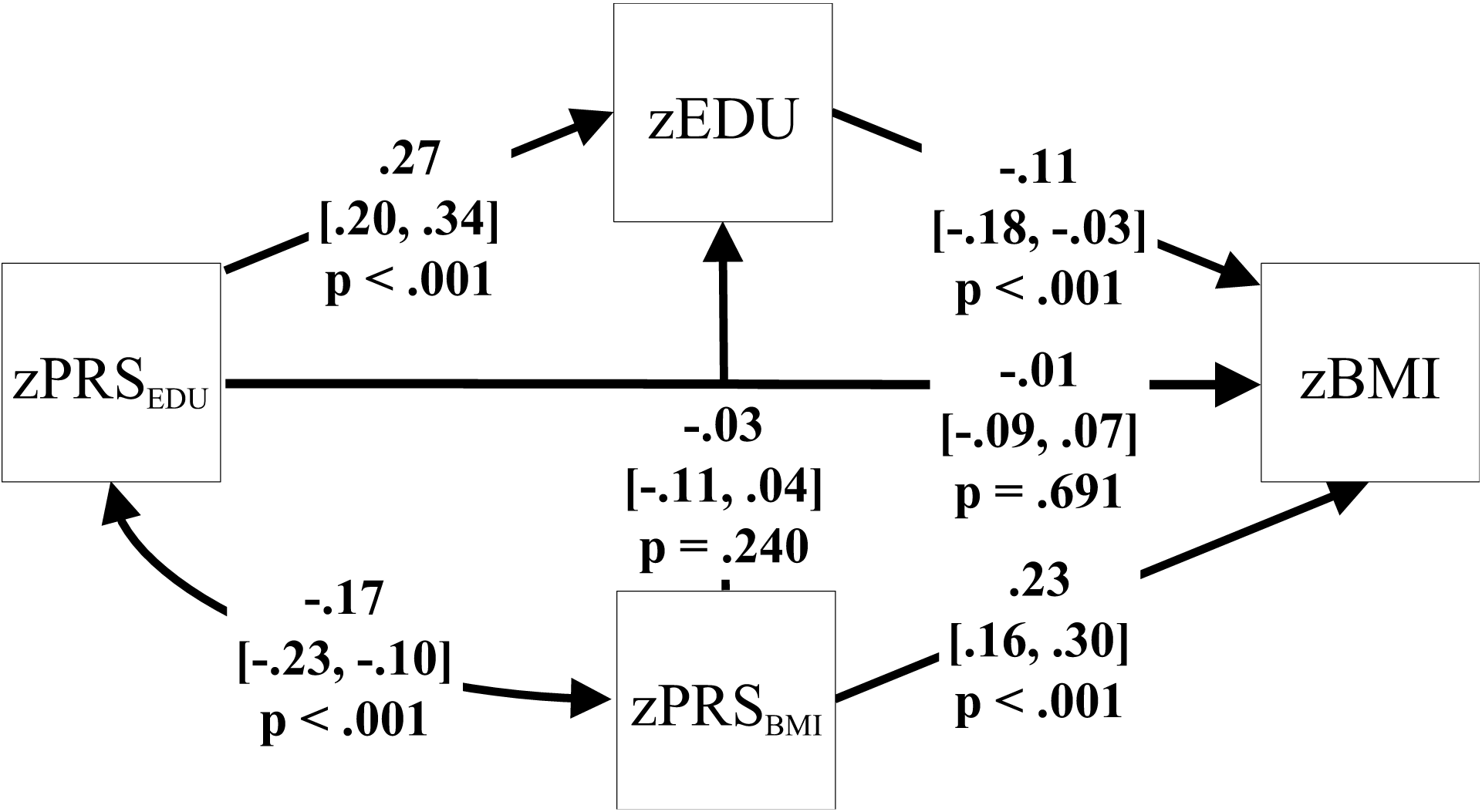
Results of a Genetic Path Analysis of Educational Attainment and Body Mass Index Using Multiple Polygenic Scores. **Notes.** The double-headed arrow represents a correlation. Single-headed arrows represent regressions. All focal variables were standardized (M = 0, SD = 1). Therefore, coefficents are intrepetted as the predicted standard deviation increase in BMI given a standard deviation increase in polygenic risk or education. 99% bias-corrected bootstrapped confidence intervals are reported below parameter estimates. p = probability of the observed data if the null hypothesis is true (i.e. β = 0). All focal variables are regressed on age, sex, and PCs, but these pathways are omitted to ease visualization. See Table 2 for the effects of exogenous covariates.

**Figure 3.**
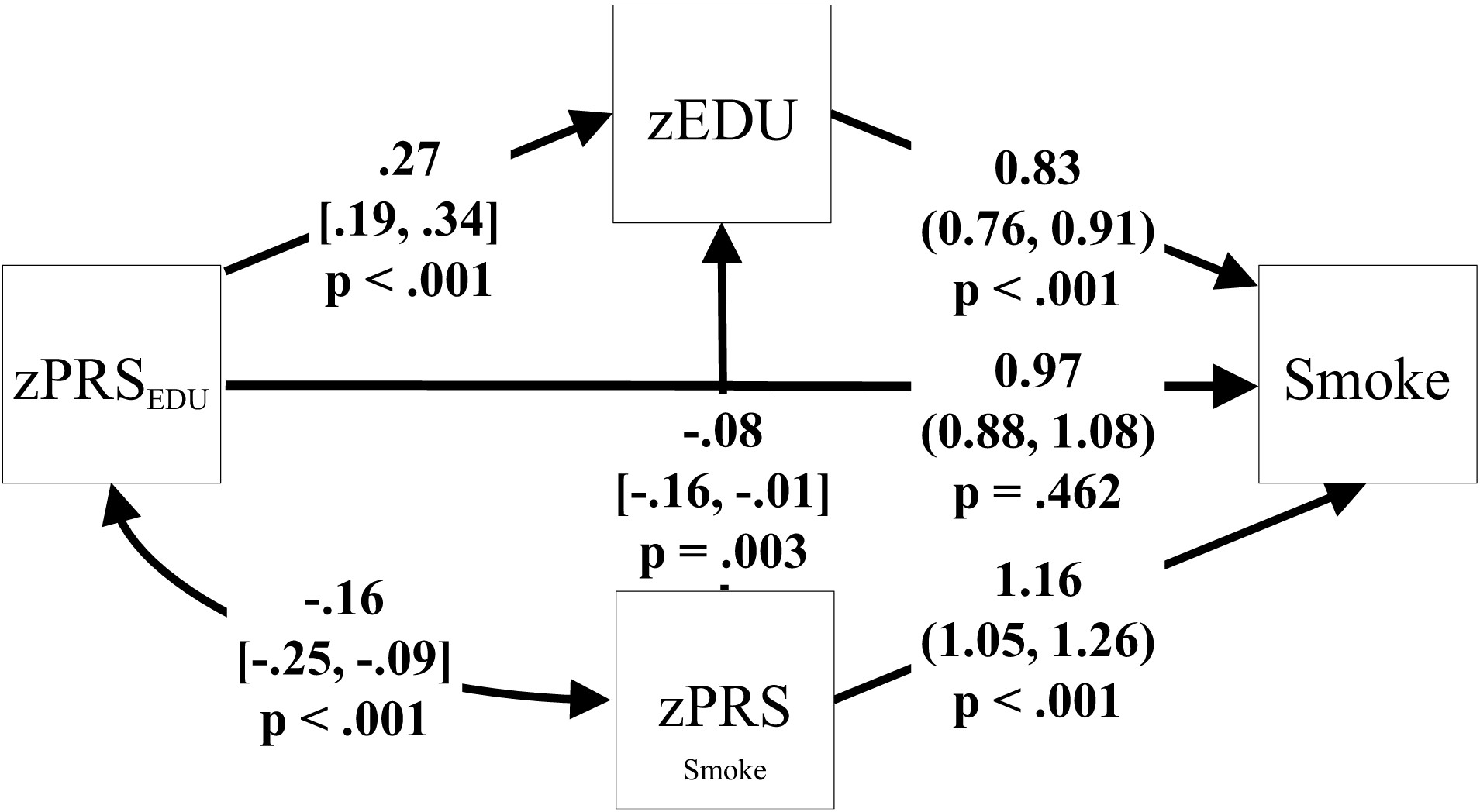
Results of a Genetic Path Analysis of Educational Attainment and Smoking Initiation Using Multiple Polygenic Scores. **Notes.** The double-headed arrow represents a correlation. Single-headed arrows represent regressions. All focal variables are standardized (M = 0, SD = 1). To help ease interpretation of results, estimates for pathways to smoking initiation are reported as risk ratios, intrepetted as the increased risk of having initiated smoking given a one unit increase in the predictor (i.e. a standard deviation increase in polygenic risk or education). 99% bias-corrected bootstrapped confidence intervals for risk ratios (RR) and betas [β] are reported in parentheses and brackets, respectively. p = probability of the observed data if the null hypothesis is true (i.e. β = 0 or RR = 1). All focal variables are regressed on age, sex, and PCs, but these pathways are omitted to ease visualization. See Table 2 for the effects of exogenous covariates

Notably, after regressing educational attainment on polygenic propensity for educational attainment, the association between polygenic propensity for BMI and education attainment approached zero (β = -.03, SE = .03, p = .240). However, even after regressing educational attainment on polygenic propensity for educational attainment, polygenic propensity for smoking initiation was negatively associated with educational attainment (β = -.08, SE = .03, p = .003). This direct association between polygenic propensity for smoking initiation and educational attainment shows that the genetic instrument for educational attainment, by itself, only provided a partial control for gene-environment correlations. The regression of the exposure on polygenic risk for the exposure *and* outcome, however, provides an additional test and control for gene-environment correlations that has not traditionally been implemented in Mendelian randomization studies. Nevertheless, even after estimating pleiotropy and polygenic propensity for the exposure *and* outcome, there was still a protective association of educational attainment on BMI (β = -.11, SE = .03, p < .001) and smoking initiation (RR = 0.83, SE = .03, p < .001). Moreover, the association between polygenic propensity for educational attainment and BMI was statistically accounted for by educational attainment (indirect effect = -.03, 99% bias-corrected bootstrapped C.I. = -.05, -.01, p = .001), as was the association between polygenic propensity for educational attainment and smoking initiation (indirect effect = -.05, 99% bias-corrected bootstrapped C.I. = -.08, -.02, p < .001).

## Discussion

The present study proposed the integration of two existing methods, genetic instrumental variable regression and path analysis, to account for pleiotropy in Mendelian randomization studies using multiple polygenic scores. The method was then evaluated using a putatively important environmental exposure and two outcomes that are of interest to clinicians and epidemiologists alike. Importantly, the present study demonstrates that education has a protective association with BMI and smoking initiation, even when controlling for potential genetic confounds via Mendelian randomization *and* pleiotropic effects using multiple polygenic scores. Moreover, for the two phenotypes examined, controls for pleiotropy were effective, such that the direct pathways from polygenic propensity for education to BMI and smoking initiation approached zero, indicating that the proposed method is capable of addressing the exclusion criteria for a sound instrumental variable. In addition, polygenic risk for smoking initiation (but not BMI) was directly associated with educational attainment, even after accounting for polygenic propensity for educational attainment. This demonstrates that, at least for some phenotypes, traditional Mendelian randomization studies provide only a partial genomic control for the environmental exposure. The method proposed and implemented in the current study, however, provides an additional test and statistical control for potential gene-environment correlations, beyond what is typically accomplished in a Mendelian randomization study.

Of course, genetic path analysis is not without limitations. For one, it can only be applied to a Mendelian randomization study for which GWAS summary statistics are available for both the exposure and outcome. In addition, although polygenic scores have become potent predictors of their respective phenotypes, especially in comparison to single genetic variants, the arrays typically included in GWASs only tag point mutations (i.e. single nucleotide polymorphisms) and do not include insertion, deletions, and copy number variants. Further, the beta weights obtained from discovery GWASs are estimated with imprecision, and, consequently, polygenic scores provide only an imperfect proxy of genetic liability. Therefore, the strength of the proposed method depends on the size and overall quality of the discovery GWASs for the exposure and outcome of interest, though the quality of the GWASs for the phenotypes examined in the present study were reasonable by contemporary standards.

In many ways, the methodological integration that was proposed and implemented in the current study is an extension or specific instantiation of genomic structural equation modeling (22). There are, however, important differences between genomic structural equation modeling and genetic path analysis as outlined in the present study. For example, genomic structural equation modeling is a technique that can be used to address a number of questions about the genetic architecture of complex phenotypes, including the search for SNPs not previously identified in a univariate GWAS. Alternatively, genetic path analysis using multiple polygenic scores was developed to address a limitation specific to Mendelian randomization studies and relies on the existence of discovery GWASs for the exposure and phenotype of interest. In addition, genomic structural equation modeling is based on genetic correlations estimated using a variant of LD-Score regression (23), and genetic path analysis relies on multiple polygenic scores to estimate genetic correlations. Genomic structural equation modeling also includes the estimation of latent variables that are not directly observed but, instead, are inferred indirectly from the data. Genetic path analysis, on the other hand, analyses associations between observed variables.

A remaining limitation to Mendelian randomization studies not addressed in the present study centers on the fact that, despite receiving a random assortment of genes from their parents, children’s genotypes depend on their parents’ genotype. Consequently, passive gene-environment correlations remain a possibility. Implementing genetic path analysis in a sample of siblings or twins would provide an additional control for this potential confound. Unfortunately, the sample analyzed in the present study did not include enough sibling-pairs to be adequately powered to fit the proposed path models to sibling-difference scores. Nevertheless, future studies may benefit from implementing genetic path analysis in larger samples of genotyped siblings with relevant exposures and outcomes measured. Finally, depicted on the top panel of Figure 1, the present study did not address potential threats to the independence criteria for a sound instrument posed by any unmeasured confounder present in a non-experimental study. Despite these limitations, the present study provides compelling evidence for a complex set of gene-environment transactions that contribute to important health-related outcomes in adulthood.

## Author Contributions

FDM developed the proposed method, conducted analyses, and drafted the manuscript. AAS & ARD performed genotype calling, imputation, and polygenic scoring. RFK contributed to the design of the study, obtained funding for the study, and supervised FDM. All authors provided critical revisions to manuscript and approved a final version.

